# Effect of temperature on mosquito olfaction

**DOI:** 10.1101/2023.04.10.535894

**Authors:** Chloé Lahondère, Clément Vinauger, Jessica E. Liaw, Kennedy K.S. Tobin, Jillian M. Joiner, Jeffrey A. Riffell

**Affiliations:** Department of Biochemistry; The Fralin Life Science Institute; The Global Change Center; Department of Entomology; Center of Emerging, Zoonotic and Arthropod-borne Pathogens, Virginia Polytechnic Institute and State University, Blacksburg, VA, 24061, USA; Department of Biology, University of Washington, Seattle, WA 98195, USA

**Keywords:** olfaction, disease vector insect, blood-feeding, *Aedes aegypti*, electroantennogram, flight arena, olfactometer, temperature

## Abstract

Mosquitoes use a wide range of cues to find a host to feed on, eventually leading to the transmission of pathogens. Among them, olfactory cues (*e.g.*, host emitted odors, including CO_2_, and skin volatiles) play a central role in mediating host seeking behaviors. While mosquito olfaction can be impacted by many factors, such as the physiological state of the insect (*e.g.*, age, reproductive state), the impact of environmental temperature on the olfactory system remains unknown. In this study, we quantified the behavioral responses of *Aedes aegypti* mosquitoes, vectors of dengue, yellow fever and Zika viruses, to host and plant related odors under different environmental temperatures.

## INTRODUCTION

Temperature is one of the most important physical factors affecting the life of insects. Because of their ectothermic nature (*i.e.*, their inability to maintain a constant body temperature, independently of the temperature of the environment), excessive heat or cold can have deleterious effects on insects’ physiology and behavior. For each insect species, its behavioral performance, such as feeding or flying, is optimal at a given temperature. Although these responses and behaviors are possible within a range of temperatures (*i.e.*, tolerance range), beyond a critical minimum and maximum point it is impossible for the insect to perform (Huey and Stevenson, 1979). Nevertheless, insects have developed strategies to avoid thermal stress and some species can rely on thermoregulation processes (Heinrich, 1993). For example, they can for adjust their body position toward the sun (May, 1976), seek shade (Herrera, 1997), use evaporative cooling of droplets of nectar (Heinrich, 1979) or urine and blood (Lahondère and Lazzari, 2012; Lazzari *et al*., 2021; Reinhold *et al*., 2021), or actively produce or release heat via counter-current heat exchangers (Heinrich 1976; Lahondère *et al*., 2017).

In mosquitoes, the deadliest animals on earth (WHO, 2019), the temperature of the environment has been shown to affect their behavior (Kirby and Lindsay, 2004; Thomson, 1938), metabolism (Clements, 1992), development (Lanciani and Le, 1995; Lyimo *et al*., 1992; Rueda *et al*., 1990), and reproduction and blood-feeding (Eldridge, 1968). Moreover, by affecting the development of the pathogens that they might carry, temperature can also affect the vectorial capacity of mosquitoes (Paaijmans *et al*., 2011; Reinhold *et al*., 2018).

In blood feeding insects, the physiological processes regulating host-seeking behavior have been well-described (*e.g.*, nutritional, and reproductive states, age, time of the day, etc. reviewed by Lazzari, 2009). By contrast, comparatively less is known about how the temperature of the environment could affect their sensory-mediated behaviors and systems, and by extension their host-seeking behavior and the transmission of the pathogens responsible for diseases. In the context of global warming and the ability of mosquitoes to invade new areas, there is an urgent need to identify the physical processes influencing mosquito behavior and biology (Lahondère and Bonizzoni, 2022). In the present study, we focused on the mosquito olfactory system – essential for host and plant seeking – and how the temperature of the environment affects mosquito responses. More specifically, we examined: (1) how temperature affects flight dynamics of tethered and free-flying mosquitoes; (2) how *Aedes aegypti*, a major vector of pathogens such as Zika, dengue or yellow fever viruses, responds to host and plant related odors under different, yet ecologically relevant, thermal conditions and (3) the possible modulation of antennal responses to odors by the environmental temperature.

## MATERIAL and METHODS

### Insects

Female *Aedes aegypti* mosquitoes (wild type, line F21 MRA-726, MR4, ATCC®, Manassas, VA, USA) were used for all experiments. The colony was kept in a climatic chamber maintained at 25±1°C, 60±10% relative humidity (RH) and under a 12-12 hr light-dark cycle. Groups of 160 - 200 larvae were placed in 26x35x4cm covered pans containing tap water and were fed daily on fish food (Hikari Tropic 382 First Bites - Petco, San Diego, CA, USA). Groups of same day pupae (both males and females) were then isolated in 16 Oz containers (Mosquito Breeder Jar, Bioquip Products, Rancho Dominguez, CA, USA) until emergence. Adults were then transferred into mating cages (BioQuip Products, Rancho Dominguez, CA, USA) and maintained on 10% sucrose. An artificial feeder (D.E. Lillie Glassblowers, Atlanta, GA, USA; 2.5 cm internal diameter) filled with heparinized bovine blood (Lampire Biological Laboratories, Pipersville, PA, USA) placed on the top of the cage and heated at 37°C using a water-bath circulation was used to feed mosquitoes weekly.

For the experiments, groups of 120 pupae were isolated and maintained in their container for 6 days after their emergence. This gave the mosquitoes time to mate in the containers before the experiments (random dissection of females revealed that 95% of them had oocytes). Mosquitoes had access to 10% sucrose *ad libitum* but were not blood-fed before the experiments. The day the experiments were conducted, females were selected manually with forceps after placing the container for a few minutes in a climatic chamber at 10°C. Depending on the type of experiment, they were then either placed individually or in groups of 20 in 50 mL conical Falcon^TM^ tubes (Thermo Fisher Scientific, Pittsburgh, PA, USA) and covered with a piece of mesh. We gave them at least 1 hr to recover before testing them in the olfactometer. All experiments were conducted when *Ae. aegypti* mosquitoes are the most active and responsive to host related cues, either 2 hours before their subjective night or 2 hours after their subjective dawn (Trpis *et al*., 1973, Vinauger *et al*., 2014).

#### 1. Tethered flight under different thermal conditions

##### 1.1. Flight arena

The in-flight response to CO_2_ of individual female mosquitoes was quantified in an LED-based arena (*sensu* Reiser and Dickinson, 2008; Vinauger *et al*., 2018; 2019). Mosquitoes were cold anesthetized on ice and tethered by the thorax with a tungsten wire using UV-activated glue (Loctite 3104 Light Cure Adhesive, Loctite, Düsseldorf, Germany), positioning their main body axis at a 30° angle from the tether. Mosquitoes were then kept at room temperature in a closed container for an approximate 30-minute recovery period. A tethered mosquito was centered in a hovering position within an arena composed of 12 columns of 2 panels each, which were arranged into a regular dodecagon and produced a display resolution of 96 x 16 pixels (Fig. 1A; Reiser and Dickinson, 2008, Vinauger *et al*., 2018; 2019). Mosquitoes were placed directly under an infrared (IR) diode and above an optical sensor coupled to a wingbeat analyzer (JFI Electronics, University of Chicago; Götz, 1987; Lehmann and Dickinson, 1997, Reiser and Dickinson 2008). The beating wings cast a shadow onto the sensor, allowing the analyzer to track their motion and measure the amplitude and frequency of each wing stroke. Measurements were sampled at 5 kHz and acquired with a National Instrument Acquisition board (BNC −2090A, National Instruments, Austin, Texas, USA). The mosquito was centered between an air inlet and a vacuum line aligned diagonally with one another, 30° from the vertical axis. The air inlet was positioned 12 mm in front of and slightly above the mosquito’s head, targeting the antennae from an angle of 15°. The vacuum line was positioned behind the mosquito 25 mm away from the tip of the abdomen. Two different airlines independently controlled by a solenoid valve (The Lee Company, Westbrook, CT, USAREF) intersected this main air inlet, either delivering clean air or CO_2_. A visual pattern of alternating stripes of either inactive or fully lit LEDs, each 16×6 pixels in size (22.5°) was used in conjunction with the odor stimulus and set in closed-loop with the mosquito behavior to maintain their motivation to fly. Odor stimuli consisted of a 1sec pulse followed by a 60-s inter-trial-interval, after which the sequence was repeated for a total of 5 pulses. The responses of the mosquitoes were tested at 20°C (n=16), 25°C (n=14) and 30°C (n=13). A nitrogen control (n= 11) was also performed.

**Figure 1.**
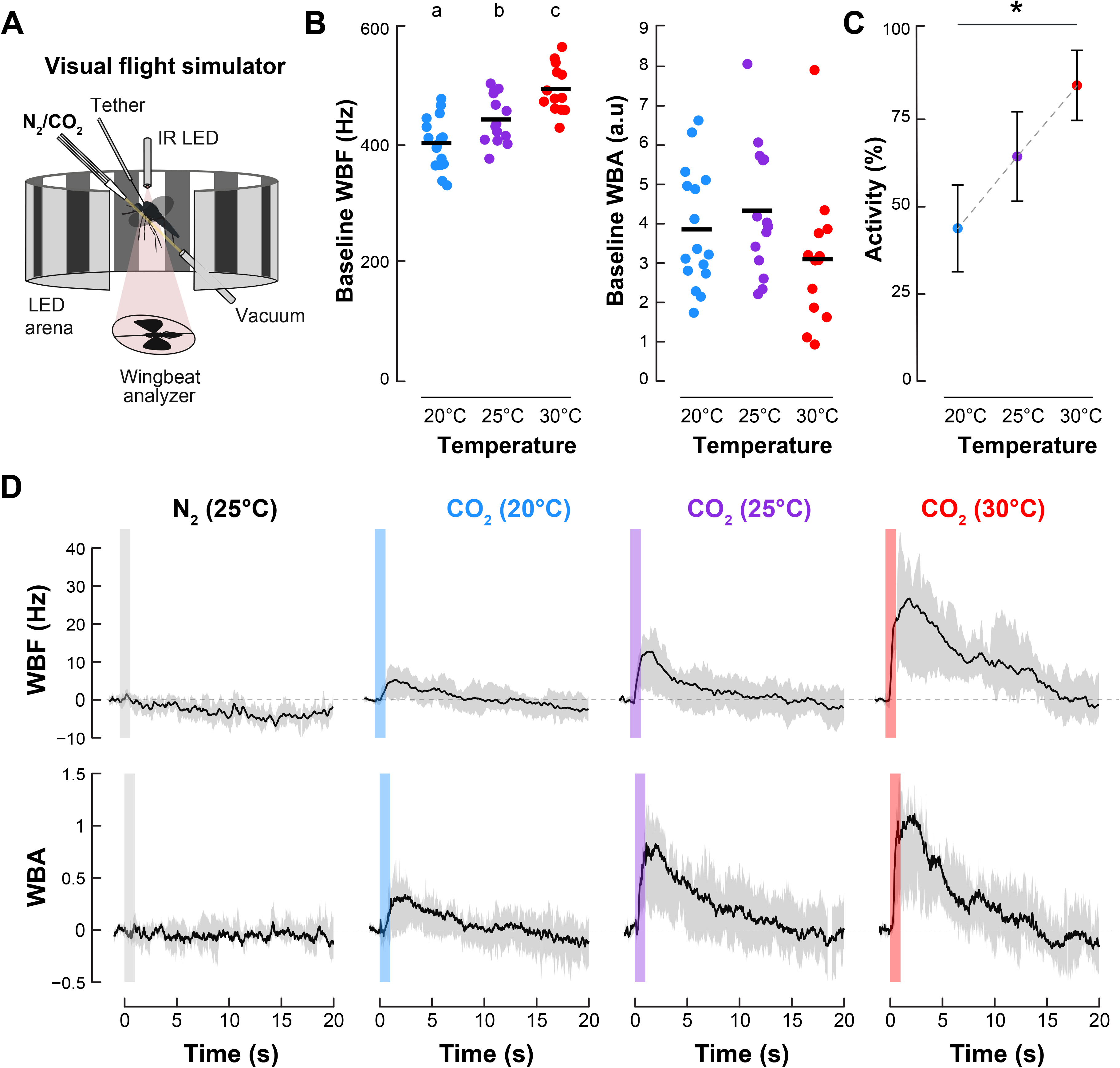
**A.** Schematic of the LED arena. The odor delivery line was connected to a thermostat-controlled conductive tubing, maintaining the main air flow at a constant temperature set at either 20, 25 or 30°C. **B.** (*Top*) trigger-averaged and baseline-subtracted wingbeat frequency (WBF), in response to a 1s long pulse of nitrogen (N_2_, grey bar, n = 11), or CO_2_ diluted into N2 at 20°C (blue bar, n = 16), 25°C (purple bar, n = 14) or 30°C (red bar, n = 13). (*Bottom*) trigger-averaged and baseline-subtracted wingbeat amplitude (WBA) in response to the same stimuli.

##### 1.2. Data analysis

Data were analyzed in R version 4.0.0. For each pulse (either CO_2_ or N_2_ control), the wingbeat frequency and amplitude were normalized by averaging the signal across a 1-s time window preceding the stimulus delivery and then subtracting this baseline value from the signal values following the pulse. Trials were discarded in which mosquitoes had frequency fluctuations greater than 5 Hz in this 1-s window or frequency changes greater than 30 Hz that did not begin within the four seconds following stimulation (as they presumably were not in response to the stimulus). The mean response for each individual was calculated from the saved trials and used as a replicate to calculate the mean response for each treatment group. This latter was calculated using the difference between the mean frequency - and amplitude - before (200 msec before stimulation) and the maxima of the signal after stimulus delivery (within a window starting at 1 s and ending 3 s after the pulse). One-tailed Student’s *t*-tests for paired samples were used to test for differences from baseline and Student’s *t*-tests for independent samples were used to test for differences between groups.

#### 2. Free flight orientation to the olfactory cue under different thermal conditions

##### 2.1. Olfactometer

To test if the ambient temperature impacts mosquito’s olfaction, we measured and compared the mosquito behavioral responses to different odors under various thermal conditions using a Y-maze Plexiglas® olfactometer. Three different temperature conditions were tested: 20°C, 25°C, and 30°C which are in the range of activity of *Aedes aegypti* (*i.e.*, 17°C to 34°C, Christophers (1960)). The olfactometer was composed of a releasing chamber connected to a tube of 30 cm long and 10 cm of diameter attached to a central box where two choice arms (39 cm long, 10 cm diameter) were placed. White cardboard was placed under and on the sides of the olfactometer to prevent the mosquitoes from being distracted by surroundings. Fans (Rosewill, Los Angeles, CA, USA) were placed at the end of the two arms of the olfactometer to generate a constant airflow (air speed approximately 40 cm.sec^–1^). To create a laminar flow, the air first entered a charcoal filter (to remove odor contaminants (C16x48, Complete Filtration Services, Greenville NC, USA)) and a series of mesh screens and honeycomb metal sheet (10 cm thick). The odor delivery system to the two choice arms was made using a charcoal-filtered air stream adjusted with flow meters equipped with needle valves. Teflon® tubing (3 mm internal diameter) conducted the air flow through each of the two 20 mL scintillation vials (Thermo Fisher Scientific, Waltham, MA, USA) containing 12 mL of either the tested odor or the control solution (*i.e.*, MilliQ water or ethanol 100%). Each tubing was connected to the corresponding choice arm of the olfactometer and placed at about 4 cm from the fans and its end located in the center of the arm. All the olfactometer experiments were conducted in a well-ventilated climatic chamber (Environmental Structures, Colorado Springs, CO, USA), 50% RH. After each experiment, the olfactometer, tubing and vials were cleaned with water and then ethanol 100% to avoid any odorant contamination between experiments. Finally, to avoid any side biases in the arms of the olfactometer, both stimulus and control vials were randomly tested on both sides for all groups.

##### 2.2. Experimental procedure

###### 2.2.1. Response to carbon dioxide in groups

The first experiment consisted in measuring the response of groups of mosquitoes to carbon dioxide (CO_2_) which is a strong activator and attractant (*i.e.*, host-seeking cue). Carbon dioxide was chemically generated using sulfuric acid (1M) and sodium carbonate (0.3M) *via* the reaction: Na2CO3 + H2SO4 → CO2 + H2O + Na2SO4. Briefly, a solution of Na2CO3 was injected at a constant rate (10 mL/h) in a jar containing 100 mL of H2SO4 using a drip-feed device (Barrozo and Lazzari, 2004). The production of CO_2_ was monitored along the experiment with a carbon dioxide detector and calibrated so the level of CO_2_ in the olfactometer was of 2000 ppm. The jar containing sulfuric acid was placed on a stir plate to help mixing the two chemicals and ensure a constant CO_2_ production. To send the odor to the olfactometer, an air pump gently injected air to the jar through tubing from the jar to the arm of the olfactometer. As a control, 100 mL of sulfuric acid was placed in a jar and the same air delivery system was used to connect to the olfactometer. Twenty female mosquitoes were released in the olfactometer and were given 30 minutes total to choose between the two arms carrying either the tested odor or the control. The number of females in each arm was then counted and the ones that stayed in the central box and main arm of the olfactometer were considered as inactive.

###### 2.2.2. Individual response to ecologically relevant odors

The individual response to different odors was then tested using the olfactometer. This allowed us to observe the behavior of each mosquito and record their flight behavior and analyze their flight trajectories. We tested seven different biologically relevant odors, both host- and plant-related: lactic acid (L-(+)-lactic, Sigma Aldrich, ≥ 98% purity - concentration: 50mM in milliQ water); Octenol (1-octen-3-ol, Sigma Aldrich, ≥ 98% purity; enantiomeric ratio: ≥99:1 (GC) - concentration: 140mM in milliQ water); Nonanol (1-nonanol, Sigma Aldrich, purum, ≥ 98.0% (GC) - concentration: 1.58mM in milliQ water), Hexanol (1-hexanol; Sigma Aldrich, > 98% purity; 92% of the Z isomer - concentration: 85mM in milliQ water), Benzaldehyde (Sigma Aldrich, purified by redistillation, ≥99.5% - concentration: 0.98M in milliQ water), DEET (N,N-diethyl-meta-toluamide, Supelco, ≥ 95% purity - concentration: 40% diluted in 100% ethanol to match commercial version of the repellent) and CO_2_ (chemically produced as previously described - concentration: 2000 ppm above ambient air levels). Hexanol and DEET are known olfactory repellents and feeding deterrents of *Ae. aegypti*, while octenol and lactic acid (here used at a concentration similar to human skin emissions (Eiras and Jepson, 1991; Cork and Park, 1996; Geier *et al*., 1996)) evoke a slight attraction preference, and nonanol is neutral. Carbon dioxide was used as a positive control and we also ran experiments with two clean air currents as a negative control and to assess that no bias (*i.e.*, innate preference for one side of the olfactometer) could be evinced. We also performed each set of experiments in duplicate, switching the odor side to control for any possible side preference. The results from the two sets of experiments were then grouped and summed. After releasing the mosquito in the olfactometer, each female was given 5 minutes to choose between the two arms. We considered that the mosquito made a choice when it crossed the entry of one of the two decision arms. If after 5 minutes the mosquito didn’t make any choice and stayed either in the central box or the main arm, then this individual was considered as inactive. A camera placed above the olfactometer allowed us to record the flight of each of the tested insects. Then, flight trajectories were recreated with video tracking using MATLAB (toolbox: DLT digitizing tools) and a mean flight velocity of the mosquitoes was calculated (as in Vinauger *et al*., 2018).

##### 2.3. Data analyses

###### 2.3.1. Comparisons of mosquito choices in the olfactometer

Binary data collected in the olfactometer were analyzed and all statistical tests were computed using R software (R Development Core Team, 2017). Comparisons were performed by means of the exact binomial test (α=0.05). For each treatment, the choice of the mosquitoes in the olfactometer was either compared to a random distribution of 50% on each arm of the maze or to the distribution of the corresponding control when appropriate. For binary data, the standard errors (SE) were calculated as in (Le, 2003):

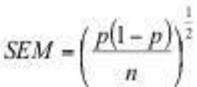

A preference index (PI) was calculated as follows: PI = [(number of females in the odor arm – number of females in the control arm) / (total number of active mosquitoes)]. A PI with a positive value indicates that the mosquitoes were attracted to the odor while a PI with a negative value indicates that mosquitoes were repelled. A PI of 0 indicates that 50% of insects chose the odor arm and 50% the control arm (adapted from Schwaerzel *et al*., 2003).

###### 2.3.2. Comparisons of mosquito velocity

Flight patterns were recreated with video tracking using MATLAB (toolbox: DLT digitizing tools) and a mean flight velocity of the mosquitoes was calculated. A Two-way ANOVA was performed to evaluate the impact of both temperature and odor on the mosquitoes’ flight velocity.

#### 3. Electroantennography (EAG)

##### 3.1. Mosquito head preparation

Electroantennograms were performed following the procedure by Lahondère (2021). Briefly, the head of the mosquitoes was excised and one segment of the tip of each antenna was cut off with fine scissors under a binocular microscope (Carl Zeiss, Oberkochen, Germany). The head was then mounted on an electrode made of a silver wire 0.01” (A-M Systems, Carlsbord, WA, USA) and a borosilicate pulled capillary (0.78 mm I.D. Sutter Instrument Company, Novato, CA, USA) filled with a 1:3 mix of saline solution (Beyenbach and Masia, 2002) and electrode gel (Parker Laboratories, Fairfield, NJ, USA) to limit desiccation during the experiment. The head was mounted by the neck on the reference electrode. The preparation was then moved to the EAG setup and using a micromanipulator (Narishige, Japan) the tips of the antennae were inserted, under the microscope (Optiphot-2, Nikon, Tokyo, Japan) into a recording electrode, identical to the reference electrode. The mounted head was oriented at 90° from the main airline which was carrying clean air (Praxair, Danbury, CT, USA) and volatiles from the syringe to the preparation for the whole duration of the experiment. Two pulses of a duration of 2.5 s and separated by 10 s were delivered to the mosquito head preparation by switching a solenoid valve controlled by a custom MATLAB script. All the chemicals tested in the olfactometer (except for CO_2_, which is detected by the mosquito palps) were used for these experiments. Pulses of clean air were used as a control as well as 100% ethanol (solvent of DEET).

##### 3.2. Data acquisition and analyses

Electrophysiological signals were amplified 100X and filtered (0.1-500 Hz) (A-M Systems Model 1800, Sequim, WA, USA), recorded and digitized at 20 Hz using WinEDR software (Strathclyde Electrophysiology Software, Glasgow, UK) and a BNC-2090A analog-to-digital board (National Instruments, Austin, TX, USA) on a personal computer. A Humbug noise eliminator (Quest Scientific, Vancouver, Canada) was used to eliminate electrical noise. The voltage pulse delivered to the solenoid valve was recorded simultaneously, on a separate channel, and a custom-written R script was used to detect the onset of the odor pulses to trigger averaging responses to each stimulus. The electrophysiological signal collected was further filtered (Butterworth digital filter set as a low pass with a Nyquist frequency of 0.99; R package *signal*). Amplitude of the EAG response was determined as the difference between the mean baseline signal in a 2 s window immediately preceding the onset of the odor pulse, and largest voltage deflection in a 5 s window following the onset of the odor pulse. A generalized linear model (glm) was fitted to the data, as: *EAG Response ∼ Temperature*odor type*. As a post-hoc analysis, general linear hypotheses and multiple comparisons were tested with the *glht* of the *multcomp* package in R.

## RESULTS

### 1. Flight arena

To examine how temperature modulates flight kinematics, mosquitoes were tethered by the thorax and maintained in an air flow at different temperatures in a LED arena (Fig. 1A). First, we found that the average baseline wingbeat frequency was lower at 20°C (408.8 ± 11.2 Hz; mean±SEM) than at 25°C (449.3 ± 11 Hz) and 30°C (501.2 ± 11.5 Hz) (Student’s *t*-tests, *p* < 0.01) (Fig. 1B). Regarding the amplitude, we found that the baseline was higher at 25°C (4.39 ± 0.45 a.u.) than 20°C (3.91 ± 0.38 a.u.) and 30°C (3.14 ± 0.50 a.u.) (Student’s *t*-tests, all *p* < 0.01) (Fig. 1B). We also found that with increasing temperature, higher the numbers of mosquitoes responded (20°C: 43%; 25°C: 64% and 30°C: 84%; Binomial Exact tests: 20-25: *p* = 0.08; 25-30: *p* = 0.05; 20-30: *p* = 0.0002) (Fig. 1C). As a control for a potential mechanical perturbation induced by the onset of the odor delivery, we stimulated mosquitoes with a pulse of the carrier gas (N_2_) at the same flow rate. Mosquitoes did not change their wingbeat frequency (Student’s *t*-test, t = −1.1013, df = 10, *p* = 0.2966) or wingbeat amplitude (t = 0.18055, df = 10, *p* = 0.8603) in response to the N_2_ pulse (Fig. 1D).

Next, changes in wingbeat frequency and amplitude in response to pulses of CO_2_ were measured. We found that the temperature affected mosquitoes’ response to CO_2_ and that the average increase was lower at 20°C (4.62 ± 1.28 Hz) than at 25°C (12.2 ± 4.22 Hz) and 30°C (22.22 ± 5.06 Hz) (20-25: t = −1.7178, df = 15.396, *p* = 0.05; 25-30: t = −1.5206, df = 23.841, *p* = 0.07; 20-30: t = −3.3693, df = 13.543, *p* = 0.002) (Fig. 1D). Moreover, the average changes in amplitude in response to a pulse of CO_2_ gradually increased with increasing temperatures (20°C: 0.28 ± 0.11 a.u., 25°C: 0.73 ± 0.19 a.u. and 30°C: 0.98 ± 0.29 a.u; 20-25: t = −2.0295, df = 20.799, *p* = 0.03; 25-30: t = −0.7196, df = 21.229, *p* = 0.24; 20-30: t = −2.2695, df = 15.431, *p* = 0.02) (Fig. 1D). The intensity of the responses to the CO_2_ pulse was higher at 30°C than at 20°C (Student’s *t*-tests, *p* < 0.01).

### 2. Olfactometer

#### Response to carbon dioxide in groups

To test whether the temperature of the environment could affect the mosquito’s olfactory behaviors, we first measured their response to carbon dioxide, an important cue used by the insects for host-seeking, in the olfactometer under different thermal conditions. We found that mosquito groups were more attracted by CO_2_ when tested at 30°C compared to 20°C and 25°C (binomial tests, p<0.05) (Fig. 2A). Moreover, the proportion of mosquitoes that made a choice in the olfactometer was also higher at higher temperatures (5.4% at 20°C, 45.3% at 25°C and 64.8% at 30°C; Chi2, p<0.05) (Fig. 2B).

**Figure 2.**
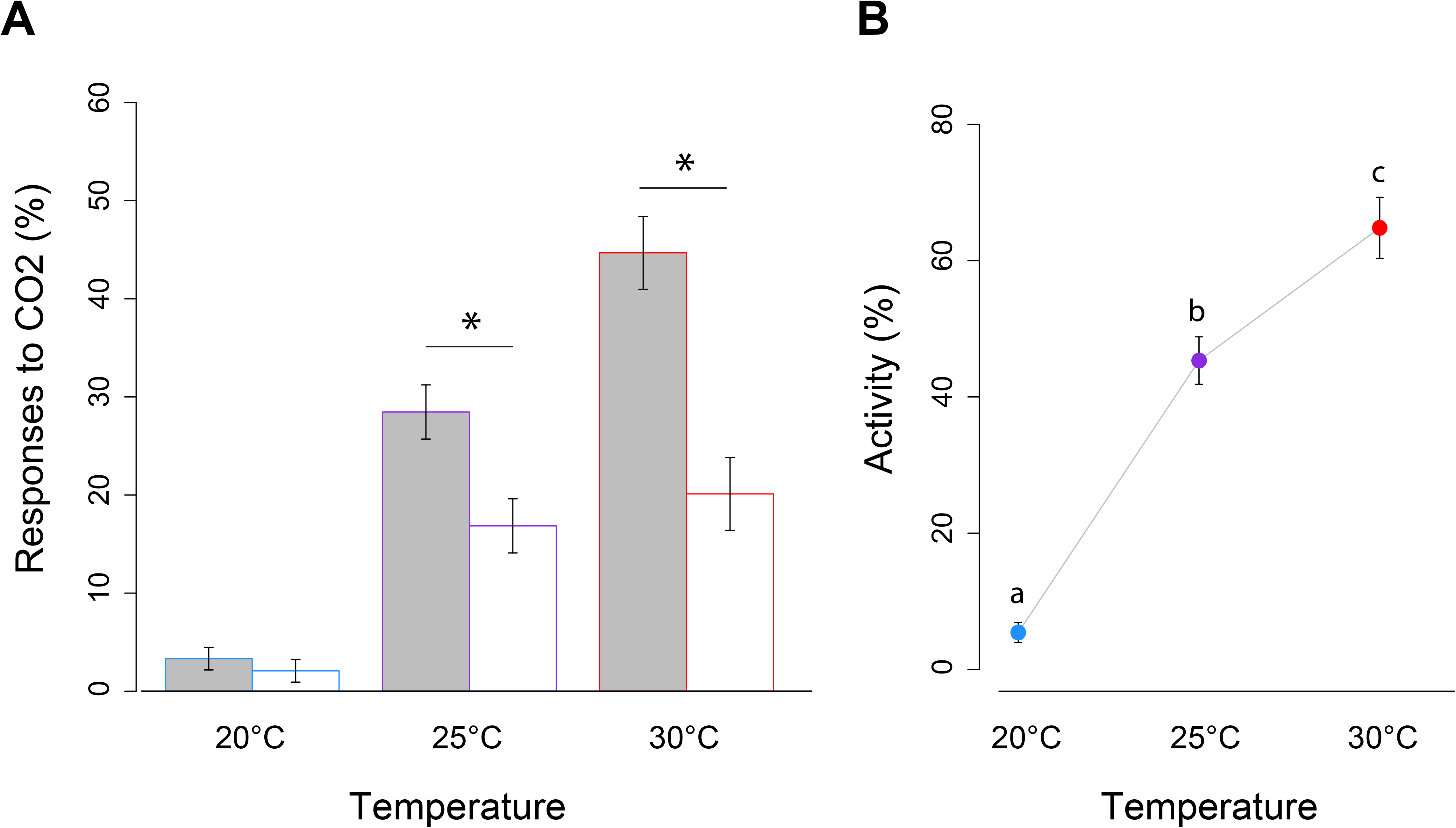
**A.** Responses of female mosquitoes to carbon dioxide tested in the y-maze olfactometer at different environmental temperatures (20°C, 25°C and 30°C). Mosquitoes were released in groups and given a choice between CO_2_ (2000 ppm) (grey bars) and clean air (white bars). Number of groups of mosquitoes tested at 20°C (N = 12, n = 240), 25°C (N = 14, n = 267) and 30°C (N = 9, n = 179). **B.** Proportion of active mosquitoes (*i.e.*, the number of mosquitoes that entered one of the two decision arms over the total number of individuals tested) at different temperatures. Asterisks indicate distributions that are significantly different from random (Binomial tests; p < 0.05). Vertical bars represent S.E.M.

#### Individual response to ecologically relevant odors

To determine whether releasing the mosquitoes in groups had any effect, and to easily track the behavior of the mosquitoes, we conducted the same experiment with individual female mosquitoes. We confirmed the results we obtained for the response to CO_2_ in groups and found that higher temperatures elicited a stronger level of attraction for the odor (binomial tests, *p* < 0.05) (Fig. 3A). We also found that hexen-1-ol, benzaldehyde and DEET significantly repelled more mosquitoes at higher temperatures (binomial tests, *p* < 0.05) (Fig. 3A). Interestingly, for some other odors such as L-(+)-lactic acid, 1-octen-3-ol and nonanol, no effect of the temperature was evinced on the mosquito choice (Fig. 3A, top panel). However, the overall level of activity was generally higher at higher temperature, independently of the odor tested (binomial tests, *p* < 0.05) (Fig. 3B).

**Figure 3.**
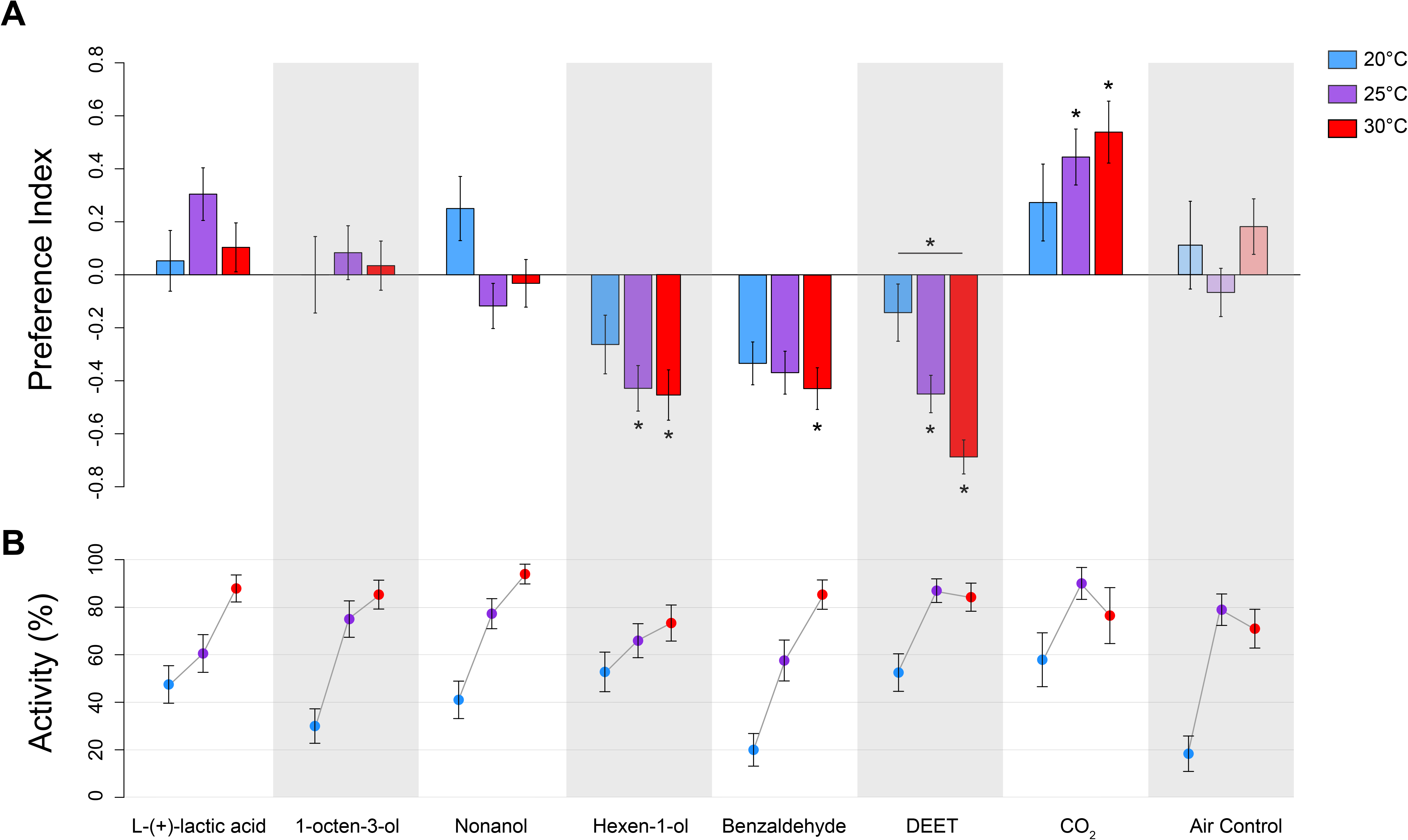
**A.** Responses of female mosquitoes to different odors, at different environmental temperatures (20°C, 25°C and 30°C). Mosquitoes were tested individually and given a choice between the test odor and a control (*i.e.*, the solvent of the odor). Mosquito choices are represented as the preference index calculated from the distribution of insects in the olfactometer. Each bar represents an experimental group tested at either 20°C (blue bars), 25°C (purple bars) or 30°C (red bars). Odor tested: L-(+) lactic acid (20°C: n = 19, 25°C: n = 23 and 30°C: n = 29), octenol (20°C: n = 12, 25°C: n = 24 and 30°C: n = 29), nonanol (20°C: n = 16, 25°C: n = 34 and 30°C: n = 31), hexanol (20°C: n = 19, 25°C: n = 29 and 30°C: n = 29), DEET (20°C: n = 21, 25°C: n = 40 and 30°C: n = 32), CO2 (20°C: n = 11, 25°C: n = 18 and 30°C: n = 13). As a control, mosquitoes were also exposed to two clean air currents (20°C: n = 5, 25°C: n = 30 and 30°C: n = 22). **B.** Proportion of active mosquitoes (*i.e.*, the number of mosquitoes that entered one of the two decision arms over the total number of individuals tested) at different temperatures. Odor tested: L-(+) lactic acid (20°C: N = 40, 25°C: N = 38 and 30°C: N =33), octenol (20°C: N = 40, 25°C: N = 32 and 30°C: N = 34), nonanol (20°C: N = 39, 25°C: N = 44 and 30°C: N = 33), hexanol (20°C: N = 36, 25°C: n = 44 and 30°C: N = 34), DEET (20°C: N = 40, 25°C: N = 46 and 30°C: N = 38), CO2 (20°C: N = 19, 25°C: N = 20 and 30°C: N = 13). A clean air control where mosquitoes are exposed to two clean air currents was also performed (20°C: N = 27, 25°C: N = 38 and 30°C: N = 31). Asterisks indicate distributions that are significantly different from random (Binomial tests; p < 0.05). Vertical bars represent S.E.M.

#### Comparisons of mosquito velocity

We then compared the flight velocity of the mosquitoes and found that temperature affected it for some odors (*e.g.*, benzaldehyde, hexen-1-ol), *i.e.*, mosquitoes flew faster at higher temperatures (Benzaldehyde: 20°C = 13 cm.s^−1^; 25°C = 19. 1 cm.s^−1^; 30°C = 18.7 cm.s^−1^; Hexen-1-ol: 20°C = 14.5 cm.s^−1^; 25°C = 14.5 cm.s^−1^; 30°C = 21.9 cm.s^−1^;), but not for others (*e.g.*, DEET, nonanol) (DEET: 20°C = 19.3 cm.s^−1^; 25°C = 16.1 cm.s^−1^; 30°C = 18.7 cm.s^−1^; Nonanol: 20°C = 15.4 cm.s^−1^; 25°C = 18.1 cm.s^−1^; 30°C = 16.1 cm.s^−1^) (Student’s *t*-tests, *p* < 0.05, ANOVA, Table 1) (Fig. 4A, 4B).

**Table 1.** Results of the analysis of variance (two-way ANOVA with interaction) on the mean flight velocity of female mosquitoes under different thermal conditions and tested for their responses to different odors. Df: degree of freedom. Sum sq: sum square, Mean Sq: mean square, F value: Mean square / Residual, Pr(>F): p-value.

| Source of variation | Df | Sum Sq | Mean Sq | F value | Pr(>F) |
| --- | --- | --- | --- | --- | --- |
| Temperature | 2 | 185 | 92.50 | 6.423 | **0.001772 \*\*** |
| Odor | 6 | 1524 | 254.05 | 17.641 | **< 2e-16 \*\*\*** |
| **Temperature : Odor** | 12 | 573 | 47.75 | 3.316 | **0.000124 \*\*\*** |
| **Residuals** | 461 | 6639 | 14.40 |  |  |

Statistically significant differences: $P < 0$ ‘\*\*\*’, $P < 0.001$ ‘\*\*’.

**Figure 4.**
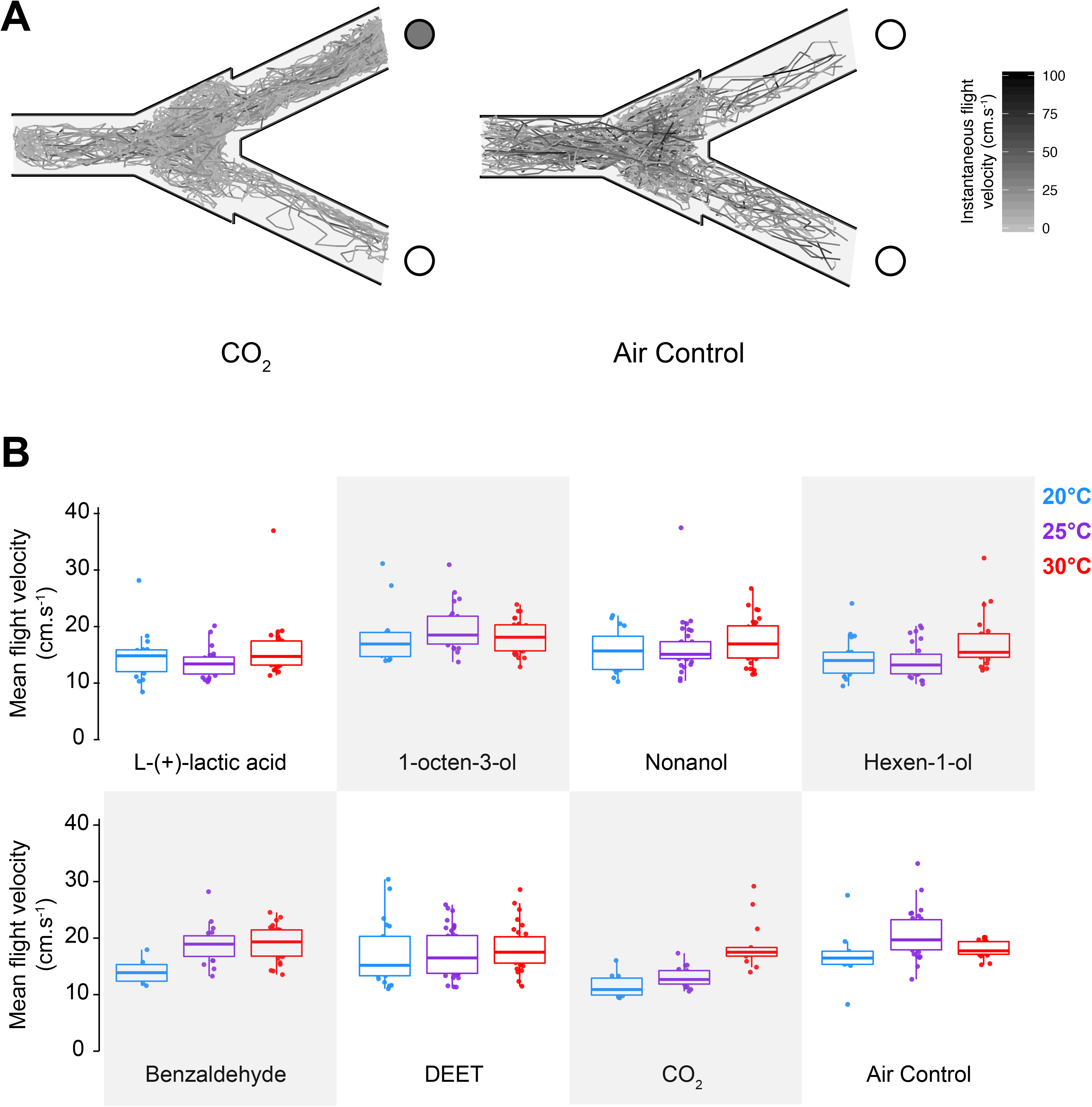
**A.** Examples of reconstructed flight trajectories obtained via video-tracking of mosquitoes in the olfactometer at 25°C. Multiple individual tracks were overlapped to obtain the present figures for mosquitoes tested for their response to CO_2_ (left; N = 18), or two control clean air currents (right; N = 30). Tracks are color coded as a function of the instantaneous flight velocity of the insect. The dark gray circle represents the arm delivering CO2, while the white circle represents the control side. **B.** Mean flight velocities (cm.s^−1^) of female mosquitoes in the olfactometer. Each boxplot represents an experimental group tested with the different odors at either 20°C (blue box plots), 25°C (purple box plots) or 30°C (red boxplots). Dots represent the mean flight velocity of individual mosquitoes; the central bar represents the median and the upper and lower bars of the plot represent the 1^st^ and 3^rd^ quartiles respectively. A two-way ANOVA was performed to compare the flight velocities under different temperature regimes according to the tested odor. Statistical results are depicted in table 1. Vertical bars represent S.E.M.

### 3. Electroantennography

Recording from the antennae of mosquitoes allowed us to quantify the peripheral olfactory sensitivity under different temperatures. Fitting a generalized linear model on the EAG data showed a significant effect of temperature (Analysis of deviance: *p* < 0.001) and of the odor identity (*p* < 0.0001) on the electrophysiological responses. Overall, responses at 25°C were significantly different from responses at 20 and 30°C (Simultaneous tests for general linear hypotheses; Multiple comparisons of means: Tukey Contrasts; 20°C vs 25°C: *p* < 0.001; 25°C vs 30°C: *p* = 0.003). However, responses at 20°C and 30°C were overall not significantly different from each other (simultaneous tests for general linear hypotheses; 20°C vs 30°C: *p* = 0.73) (Fig. 5).

**Figure 5.**
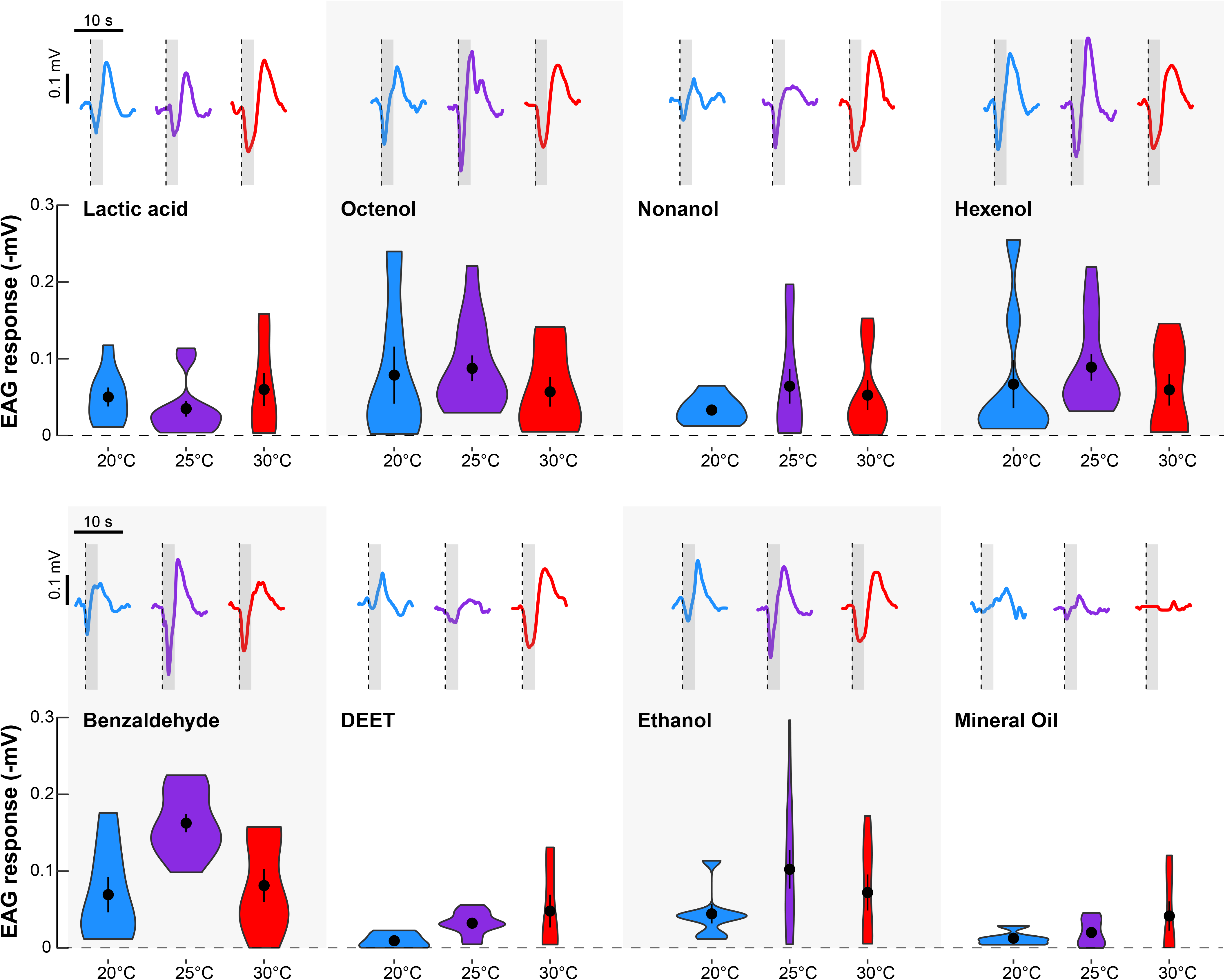
Violin plots of the EAG responses (- mV) to the different chemicals tested under different thermal conditions. The central black dot indicates the mean, and the vertical bar represents S.E.M. Above each violin plot is a representative EAG response. The grey bar indicates the odor pulse. Note the typical “V” shape of the EAG responses as well as the absence of antennal response to the solvent (no odor) control, mineral oil.

No significant effect of the interaction between temperature and the odor identity was found (Analysis of deviance: *p* = 0.078), although we observed a trend for some odors (*e.g.*, octenol, nonanol, hexanol, benzaldehyde, ethanol), but not others (*e.g.*, lactic acid, DEET), to display an optimum in EAG response at 25°C (Fig. 5).

## DISCUSSION

The present study provides clear evidence that temperature modulates mosquito olfactory behavior in both tethered preparations and free flight. In addition, our electrophysiological results show that the effect of ambient temperature on mosquito behavior is mediated by a modulation of the antennal sensitivity to odors. The approach employed here allowed us to discriminate between effects on the overall activity of mosquitoes and effects on the oriented response to odorants. Indeed, the proportion of responsive mosquitoes tended to be higher when the temperature increased in both the flight arena and the olfactometer, but the response of the mosquitoes was modulated in an odor-dependent manner. While a larger proportion of mosquitoes made a choice at higher temperature in free flight, this response remained neutral for several chemicals at the three tested temperatures. Interestingly, a stronger level of attraction (*i.e.*, CO_2_) or aversion (*i.e.*, benzaldehyde, DEET, and hexen-1-ol) were noted for several chemicals, indicating that temperature affects mosquito olfactory choices as well as their flight kinematics. In the arena, we noted that baseline wing beat frequency proportionally increased with temperature, while the amplitude of wing motion was maximal at 25°C. It is well known that ambient temperature (T*_a_*) affects insect flight (*e.g.*, Heinrich, 1993, Reinhold *et al*., 2018) and that the flight force generated by insect wings is affected by changes in stroke amplitude and frequency (Lehmann and Dickinson, 1997). Using a mosquito flight mill, Rowlet and Graham (1968) found that 21°C is the optimal flight temperature in terms of flight distance and duration but indicated that the maximum flight speed occurred between 27°C and 32°C. Christophers (1960) also highlighted that flight frequency increases with T*_a_*, reporting 367 beat/s at 18 °C vs. 427 beat/s at 25 °C in *Ae. aegypti*. Gopfert *et al*. (1999) and Arthur *et al*. (2014) recorded higher average frequencies, ranging from 445 Hz to 510 Hz at medium temperature (23°C). More recently, Villarreal *et al*. (2017) showed an increase of 8 - 13 Hz in wing beat frequency for every 1°C gain in *Ae. aegypti*. Temperature also affected mosquito flight velocity in the olfactometer, which increased in response to some odors but not others. In our olfactometer experiments, in both group and individual assays, we found that more mosquitoes made a choice towards the arm delivering the CO_2_ at 30°C than 20°C. It is worth mentioning that more mosquitoes were active at higher temperatures but the nature of their choice (*i.e.*, the preference for the test odor arm among the mosquitoes that chose one of the two arms) also changed at higher temperatures.

Temperature is known to affect neuronal activity in both vertebrates and invertebrates (Martin *et al*., 2011). In the present study, we assessed whether the peripheral detection of chemicals by the antennae was affected by T*_a_*. We found that responses of the antennae varied with T*_a_* but were also odor-dependent, reflecting effects observed in the olfactometer. Our data indicate an optimum at 25°C for antennal sensitivity. Martin *et al*. (2011) found that exposure to a higher T*_a_* led to an increase in the EAG response amplitude. Moreover, using single sensillum recordings (SSRs), they noted that the ORNs were directly impacted by T*_a_*. It is worth mentioning that although sensitivity peaks at 25°C in the EAG, some odors still show higher behavioral attraction levels at 30°C which could be due to an effect of the temperature at the central level, *e.g.*, at the level of the antennal lobes. Moreover, a higher antennal sensitivity did not necessarily correlate with a higher behavioral response, as expected for odors presented alone. This further supports the hypothesis of an additional layer of modulation of T*_a_*, at the level of central olfactory processing. Future experiments including antennal lobe recordings will reveal whether T*_a_* has an impact on signal processing in the brain. Moreover, testing the impact on temperature on ORs sensitivity in mosquitoes could also help determine why T*_a_* affects behavioral responses to certain odors but not others.

*Aedes aegypti* is a major disease vector and an invasive species which global distribution is expected to shift because of global warming (Reinhold *et al*., 2018). Increased average daily temperatures will impact the mosquito’s general activity, including flight, as well host-seeking and biting behavior, which could have major consequences for public health due to increased pathogen transmissions. It is thus critical to understand how T*_a_* affects mosquito behavior and in particular olfactory behaviors which are central to host seeking. Finally, as we are facing challenges in mosquito control due, in part, to insecticide resistance, a better understanding of how environmental factors affect mosquito behaviors can provide important knowledge to trap and bait design. The present data provide critical insights on how T*_a_* affects both the mosquito olfactory system and its flight kinematics which can be integrated in traps relying on olfactory baits as well as acoustic traps (Andrés *et al*., 2020).

## Acknowledgments

We thank Binh Nguyen and Mikayla Kiyokawa for taking care of the mosquito colony. We would like to thank Raymond Huey for helpful discussions. We are indebted to *The Journal of Experimental Biology Travelling Fellowship* awarded to Chloé Lahondère. Jessica Liaw and Kennedy Tobin were supported by *The Mary Gates Fellowship*. We acknowledge the support of the Air Force Office of Scientific Research under grants FA9550-16-1-0167 (JAR), and FA9550-21-1-0101 (JAR); the National Science Foundation under grant 2121935 (JAR); and the National Institutes of Health under grant R01AI148300 (JAR).

